# Induction of Reproductive Behaviors by Exogenous Hormones in Captive Southern Rocky Mountain Boreal Toads, *Anaxyrus boreas boreas*

**DOI:** 10.1101/131763

**Authors:** Natalie Emma Calatayud, Amanda Kathryn Mullen, Cecilia Jane Langhorne

**Affiliations:** Mississippi State University, Department of Biochemistry and Molecular Biology, Entomology and Plant Pathology, Starkville, MS 39769, USA

**Keywords:** Amplexus, Gonadotropin releasing hormone (GnRH), Human chorionic gonadotropin (hCG), Oviposit hormone dose (OvD), Post hormone–treatment (PT), Southern Rocky Mountain boreal toad (SRM)

## Abstract

Loss of reproductive viability, physiologically and/or behaviorally, can have profound effects on the fitness of a captive population and conservation efforts. The southern rocky mountain (SRM) population of the boreal toad has declined over the past 35 years, making captive breeding necessary to protect and augment the species in the wild. In recent years, a notable reduction in the incidence of amplexus and viable offspring from the captive breeding population has been observed. Hormone treatment protocols to stimulate gamete release in males and females are established in this species and *in vitro* fertilization has been performed successfully. However, successful hormone stimulation of reproductive behaviors and natural fertilization has not been well documented. During the breeding season of 2012, 24 males and 24 female toads were selected from a population of over 600 captive animals. Both sexes were treated with Human chorionic gonadotropin (hCG) and Gonadotropin Releasing Hormone (GnRH) or phosphate buffered saline (PBS). Females were primed twice with 3.7IU/g hCG and then injected with an ovulatory dose (OvD) of 13.5 IU/ g BW (Body weight) hCG and 0.4 μg/ g BW GnRH. Males were injected a single time with 10 IU/g BW hCG and 0.4 μg/ g BW GnRH, 12 h after females received their OvD. In 2013, knowing the approximate time when females oviposited after hormone treatments, we tested the best time to induce amplexus and spermiation. Males were divided into 4 groups and injected at 4 different times: (a) 12 h before females OvD; (b) at the same time as OvD; (c) 12 h after OvD; (d) control injected with PBS. Results from 2012 indicated that oviposition was solely dependent on females receiving hormone treatments not males. However, in 2013 we found that the duration of amplexus significantly influenced oviposition (*P*>0.05), and males injected 12 h prior to females spent more time in amplexus than males injected at the same time or 12 h after the females received hormones. Promoting reproductive behaviors and synchronizing gamete deposition continues to be imprecise and may require more than exogenous hormones. The complexity of promoting breeding behaviors may require a closer assessment of the captive environment.

---

LOW REPRODUCTIVE capacity is not uncommon in captive amphibian populations and has recently been observed in a captive population of the Southern Rocky Mountain boreal toad (SRM), *Anaxyrus boreas boreas.* Human chorionic gonadotropin (hCG) and gonadotropin releasing hormone (GnRH) can induce amplexus and spermiation (Roth et al. 2010) in male boreal toads, abdominal contractions and oviposition in the females (Calatayud et al. 2015), and can be administered to collect gametes independently for *in vitro* fertilization. However, there is little information about when hormones should be administered to males and females to induce reproductive behaviors and synchronize gamete deposition.

In the wild, boreal toads breed after a long period of brumation in the spring or early summer, depending on elevation and the time of year when snow disappears (McGee and Kenaith 2004). Breeding begins when males amplex females, which can last for hours or days before eggs are deposited (McGee and Kenaith 2004; personal observation). In the wild, females lay between 6000–12,000 eggs per clutch and may be biennial breeders (Hammerson 1999; McGee and Kenaith 2004; Carey et al. 2005; Muth et al. 2013). Biennial breeding is a parameter often overlooked in amphibian captive breeding programs. In temperate amphibian species that inhabit high elevations, particularly for females, breeding is determined by the availability of energy reserves, the environmental conditions, and the risk breeding has on an individual’s lifetime fitness (Muths et al. 2010, 2013). Thus, it is likely that loss of reproductive behaviors in captivity is associated with inappropriate environmental stimuli and inadequate or low nutrient availability. Similar to other amphibian species, captive boreal toads of breeding age (≥6 years for females; ≥ 4 years for males) that fail to exhibit normal reproductive behaviors after artificial brumation can be treated with exogenous hormones to stimulate reproductive behaviors and promote gamete release (Johnson et al. 2002; Herbert 2004; Michael et al. 2004; Browne et al. 2006a,b; Trudeau et al. 2010; Silla et al. 2011; Kouba et al. 2009; Roth et al. 2010; Kouba et al. 2012; Calatayud et al. 2015; Theo Smith, personal observation). In the last 10 years, SRM boreal toads at the Native Aquatic Species Restoration Facility (NASRF, Alamosa, CO) were treated with hCG and luteinizing hormone releasing hormone (LHRH; or alternatively, gonadotropin releasing hormone, GnRH), when reproductive behaviors, such as amplexus and oviposition, did not occur naturally (Theo Smith, personal communication).

However, no systematic studies have addressed the efficacy of the hormone protocols, whether both males and females require hormone treatments, and how efficient treatments are in promoting and enhancing reproductive behaviors and gamete production in the SRM boreal toad.

To first explore the induction of reproductive behaviors in SRM boreal toad females and the importance of the male presence for oviposition we examined: (1) the induction of amplexus, oviposition, and tadpole production after administration of hCG and GnRH to males alone, females alone or both sexes simultaneously compared to control animals; and 2) the effect of hormone treatments on breeding when administered to males at different time periods in relation to the female’s ovulatory dose.

## MATERIALS AND METHODS

### Housing and Maintenance

Male and female boreal toads were housed together in groups according to the region from which original founders were obtained at the Native Aquatic Species Restoration Facility in Alamosa, Colorado (NASRF). NASRF toads were brumated between December and May at temperatures between 2–6°C in an EcoPro G2 1350 Liter upright refrigerated cabinet (Foster refrigerator, Corp., Hudson, New York, USA) in plastic boxes (33 × 13 × 15 cm) lined with a layer of activated carbon, moistened sand (3.81 cm deep) and moistened sphagnum moss.

The number of individuals per group ranged from 5 to 20 and often contained multiple generations from a particular region. During the active season (outside the brumation period), toads were housed in rectangular fiberglass tanks (121 × 60 × 30 cm) tilted at a 20° angle to allow constant drainage of free–flowing groundwater. Crickets and mealworms were gut–loaded with Bug Burger®, Hydro–load® water replacement gut– load (Allen Repashy’s®, La Jolla, California, USA) and fresh carrots prior to being fed to the toads. Toads were fed live prey 3 times per week.

Before reintroducing animals to specific locations in the wild, individuals from the captive colony were selected for breeding according to age (≥ 4 years for males and ≥ 6 years for females) and county of origin, and were interbred according to a particular pedigree. Therefore, male:female ratios were dissimilar as were the numbers of animals per experimental group. In addition, females were only selected for breeding if they had not oviposited the previous year or had not been used for breeding in 2–4 years. The water temperature in the tanks during the breeding season ranged from 15–18.3 °C. Toads were exposed to natural rather than artificial lighting. Egg clutches were removed from the breeding tank 24 h after oviposition was complete and were transported to a separate rearing facility.

### Hormones

Two exogenous hormones were used during this study: an LHRH analog ([des– Gly10], D–Ala6 ethylamide acetate cat#: L4513; Sigma–Aldrich, St. Louis Missouri, USA) and human chorionic gonadotropin (hCG; cat# C1063; Sigma–Aldrich, St. Louis, Missouri, USA). hCG was reconstituted in sterile PBS at 2,500 IU or 5,000 IU per milliliter and GnRH solution was prepared at 0.4μg / 5 μL. Priming and ovulatory doses are described below. All doses were administered per gram body weight (g / BW). GnRH solutions were stored in 1mL aliquots at –20°C hCG and thawed on the day of injection. Hormones were injected intra–peritoneal (IP) using a 27–gauge needle. Females were treated with two priming doses of hCG (3.7 IU/g BW) 72 h apart and the final OvD of hCG (13.5 IU/g) and GnRH (0.4 μg/g) combined was administered 24 h after the second priming dose (Calatayud et al. 2015). Males received a combination dose of hCG (10 IU/g) (Langhorne, CJ personal communication) and GnRH (0.4 μg/g) (Trudeau et al. 2010). Control males and females received injections of Phosphate buffered saline (PBS) alone (200 μL).

### Experiment 1

In 2012, an experiment was designed to test the efficacy of hormonal induction of amplexus in males, and oviposition in females. Reproductive responses in toads after administration of hCG and GnRH were compared to control animals injected with PBS. Females (*n* = 24) were randomly assigned to two groups, a hormone treatment or a control group, of 12 females each.

During this experiment, the number of males’ amplexing and the number of females depositing eggs was recorded. The number of eggs was counted whether an amplexed or an un–amplexed female deposited them. The experiment was terminated 7 days after the OvD if females had failed to oviposit. The experiment was also terminated if a male and female in amplexus also failed to produce an egg clutch after 7 days. This termination time point was based on previous observations that non–amplexed females oviposit between 72–96 h after their OvD (Calatayud et al. 2015) and amplexed females between 24–48 h post OvD (personal observation).

### Experiment 2

In 2013, a second experiment examined the administration of hormone treatments when administered to males at 3 different times, (a) hormone administration 12 h before the female OvD (*n* = 7 males); (b) concurrently (at 0 h) with the female’s OvD (*n* = 6 males); (c) 12 h post–OvD (*n* = 4 males); (d) controls (PBS) (*n* = 5 males). Five control males were housed as follows: 2 male in groups A, 1 male in group C and 2 control males in group B. In each group control males were injected with PBS at the same time as the other males were injected in their respective groups. The uneven numbers of males in groups A–C were the result of an unbalanced ratio of males to females in the colony, and the particular combinations of pedigrees that were recommended for breeding. Therefore, 21 males were given access to 31 females and once a male had selected a female, unpaired females were removed from the tank and housed separately. In this experiment, all 31 females were treated with hormones as described in the 2012 experiment.

During this study, the percentage of males’ amplexed post treatment (PT), the duration of amplexus, and the number of females (whether amplexed or un–amplexed) that oviposited was recorded. Males were first observed 1 h after the injection and then monitored every hour for 12 h until midnight. Observations were resumed at 6am and the percentage of males observed amplexing each morning was determined. Spermic urine from amplexed males was collected by catheterization as described in Kouba et al. (2012). However, to avoid over handling the animals during amplexus, the presence of sperm was noted by collecting at least 5 μL of spermic urine but concentration and motility were not recorded.

Statistical analysis was carried out in R–studio (RStudio 0.99.489, © 2009–2014 RStudio, Inc., Cary, NC, USA) and the significance was set at P<0.05. Data were expressed as the mean ± standard error. Shapiro–Wilks test for normality showed that all our data sets were not normally distributed. The data were further analyzed using a Levene’s test to assess the equality of variances, which was found to be true for all data sets. The effect of hormone treatment on males and females and the effect of amplexus on mean oviposition rates were analyzed by ANOVA in relation to (1) hormone treatment of males, females, or both compared to controls, and (2) amplexus and (3) time to oviposition.

In 2013, a Shapiro–Wilks test for normality showed that all our data sets were not normally distributed. Once again, a Levene’s test was performed and the assumption of equal variances was found to be true for all data sets. The effect of hormone administration on males and the, (1) time to amplexus, (2) duration of amplexus, (3) oviposition with or without amplexus, and 4) time to oviposition in amplexed versus un– amplexed females at 3 different times were analyzed by ANOVA.

## RESULTS

### Experiment 1

In 2012, hormone treatment significantly affected the number of females that oviposited (*P* < 0.05; Table 1) but oviposition was not affected by amplexus (*P* > 0.05). Oviposition in 7 hormone–stimulated amplexed females and 5 un–amplexed females was observed compared to 1 amplexed control female. Furthermore, there was a significant difference (*P* < 0.05) in the time to oviposition by non–amplexed (72 and 96 h post–OvD injections) and amplexed females (28 to 48 h post–OvD injections).

**TABLE 1.** Shows the number of females in control and hormone treated groups, the number and percentage that oviposited, were amplexed, oviposited with or without the aid of a male (without amplexus). In 2012, no significant differences were observed between the females that oviposited after amplexus and those that oviposited without amplexus. However, there was a significant difference between the number of eggs oviposited between control and hormone treated females.

|  | Treatment |  | *P*-value |
| --- | --- | --- | --- |
|  | Control | Hormone |  |
| Total number of females | 12 | 12 |  |
| Number of amplexed females | 2 | 7 |  |
| Percentage amplexed | 17 | 58 |  |
| Number of females ovipositing | 2 | 4 |  |
| Percentage females oviposited | 17 | 33 |  |
| Total eggs produced | 3,726 | 15,987 |  |
| Mean number of eggs | 1,863 | 3,997 | *P* < 0.05 |
| Total number of eggs with male amplexus | 3,726 | 4,525 |  |
| Mean number of eggs with male amplexus | 1,863 | 2,284 | *P* > 0.05 |
| Total number of eggs without male amplexus | 0 | 11,462 |  |
| Mean number of eggs without male amplexus | 0 | 2,292 | *P* > 0.05 |

Males began amplexing 6–10 h after they were injected. Treating males with hormones did not increase the probability that they would amplex females (P > 0.05).

### Experiment 2

In 2013, experiments using hormones to induce amplexus were redesigned based on 2012 observations. In this study, males were separated into four groups and injected at different times with respect to female injections. Although hormone treatments did not have a significant effect on the induction of amplexus (*P* > 0.05) and amplexed males were not more likely to induce oviposition (*P* > 0.05), the time spent in amplexus was significantly influenced by the time at which they were treated with hormones (P < 0.05).

The average time taken for a male to amplex was 12–14 h PT. The time spent in amplexus was significantly different (26 h PT; *P* < 0.05) between males in group A (injected 12 h prior to OvD) compared to the other three groups, B (injected at the same time as OvD), C (injected 12 h after OvD) and control, (injected with PBS according to the treatment group with which they were housed A, B or C). The percentage number of males in amplexus at 12 h PT was greater in group A (57 %) than in groups B (17 %), C (25 %) and control (20 %), and remained higher than other groups after 26 h PT. At 26 h PT, group A had 57 % of males still amplexed, while group B and C had 0% males amplexed and the control group had 50 % (Table 2). Males in group A spent more consecutive hours in amplexus with a mean of 34±4.36 h before oviposition occurred, compared to males in the control group and groups B, C which had a mean of 18.38 h ±3.06 h. The total number of eggs oviposited in group A was higher than in other groups but the mean number of eggs oviposited among the groups was not significantly different (Fig. 1). Collection of spermic urine was performed to verify a complete reproductive response to hormones. Hormone–treated males showed an initial production of sperm at 3 h PT before becoming amplectic indicating a faster spermiation than behavioral response to the hormone. In three control males, sperm was first detected between 9–12 h after PBS injection, when amplexus was first detected. The presence of sperm in control males, continued to be observed in samples collected at 24 and 48 h PT the same duration as hormone–treated males.

**TABLE 2.** Male hormone treatment summary shows number of males treated with hormones in each group, the number and percentage that amplexed a female and the number of eggs that resulted from those pairings.

|  | Male number | Amplexed pairs | % Amplexed<br>pairs | Total eggs |
| --- | --- | --- | --- | --- |
| Control | 5 | 1 | 20 | 5,926 |
| Group A | 7 | 4 | 57 | 9,646 |
| Group B | 6 | 1 | 17 | 1,852 |
| Group C | 4 | 1 | 25 | 2,436 |

**FIG. 1.**
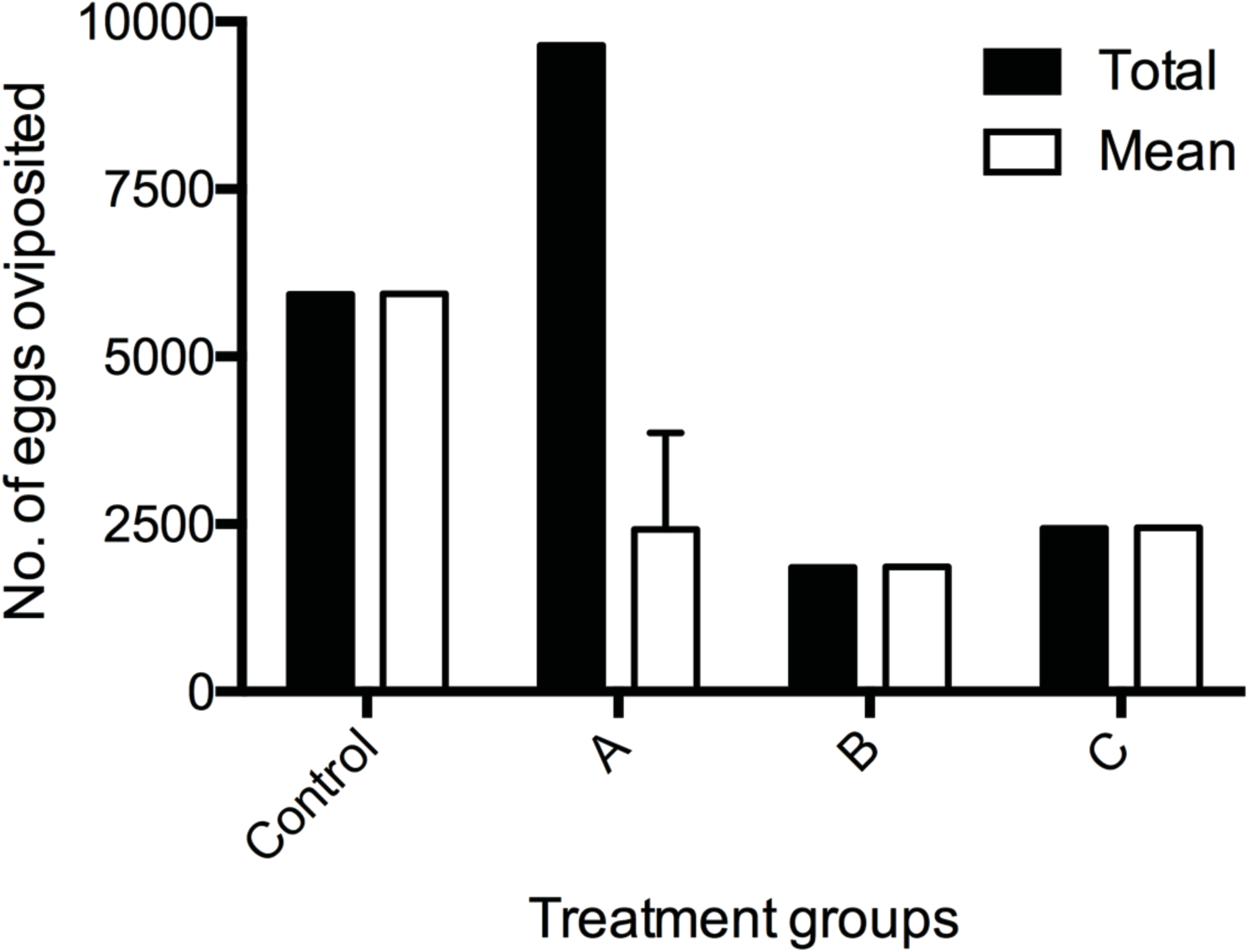
Total number of eggs deposited by amplexed pairs during the breeding season of 2013. Treatment groups reflect the differences in the time at which males were injected, since all females received their injection at the same time (with male group B). Male treatment group’s A) hormone administration 12 h before the female OvD (*n* = 7 males); group; B) concurrently (at 0 h) with the female’s OvD (*n* = 6 males); group; C) 12 h post–OvD (*n* = 4 males); D) control males (PBS) (*n* = 5 males). Although significantly more egg clutches were oviposited by pairs from group A (3 clutches) compared to groups’ B (1 clutch), C (1 clutch) and control (1 clutch), the mean number of eggs deposited per clutch was not significantly different between treatment groups.

During the 2013 experiment, no significant difference was observed between oviposition in hormone–treated amplexed and unamplexed females (Fig. 1; *P* > 0.05). The average time to oviposition for amplexed females was approximately 22.24 h (±1.36 h) PT (Fig. 2), which was shorter than the oviposition times observed for females in 2012. Additionally, three females were observed ovipositing in the absence of an amplexed male 14 h PT. Although more clutches and a larger number of eggs were oviposited by animals in group A, the mean number of eggs between groups was not different (Fig. 3). Nineteen thousand seven hundred and thirteen eggs were produced from 14 clutches in 2012 and 25, 750 eggs were produced from 10 clutches in 2013.

**FIG. 2.**
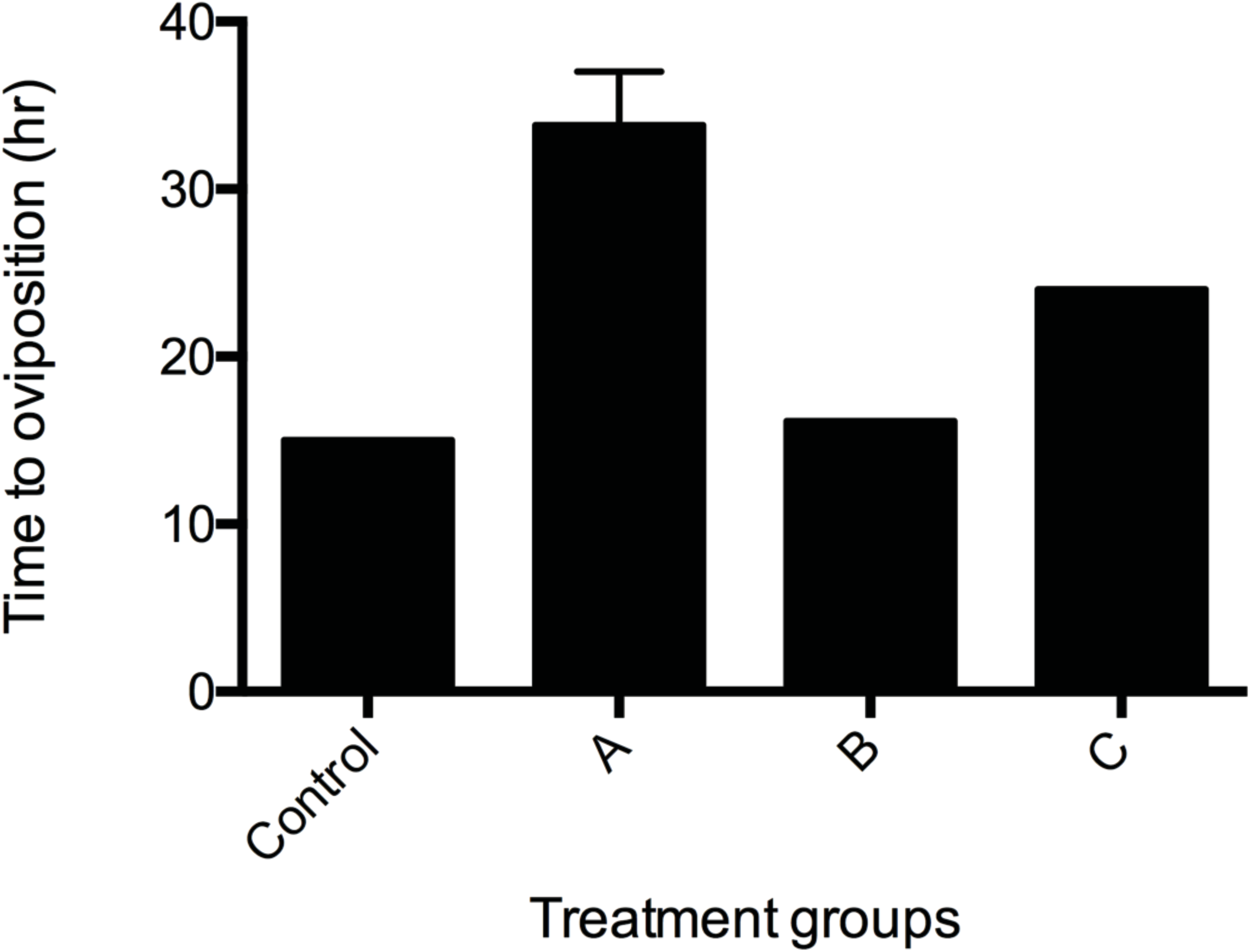
Shows the time to oviposition by amplexed pairs in 2013. The time to oviposition in groups A, B, C and control was significantly different most likely due to the time at which the females were injected. Males’ from group A were already amplexed when females housed with them received their OvD therefore, group A males spent more time amplexed before females (stimulated by hormones) were able to oviposit.

**FIG. 3.**
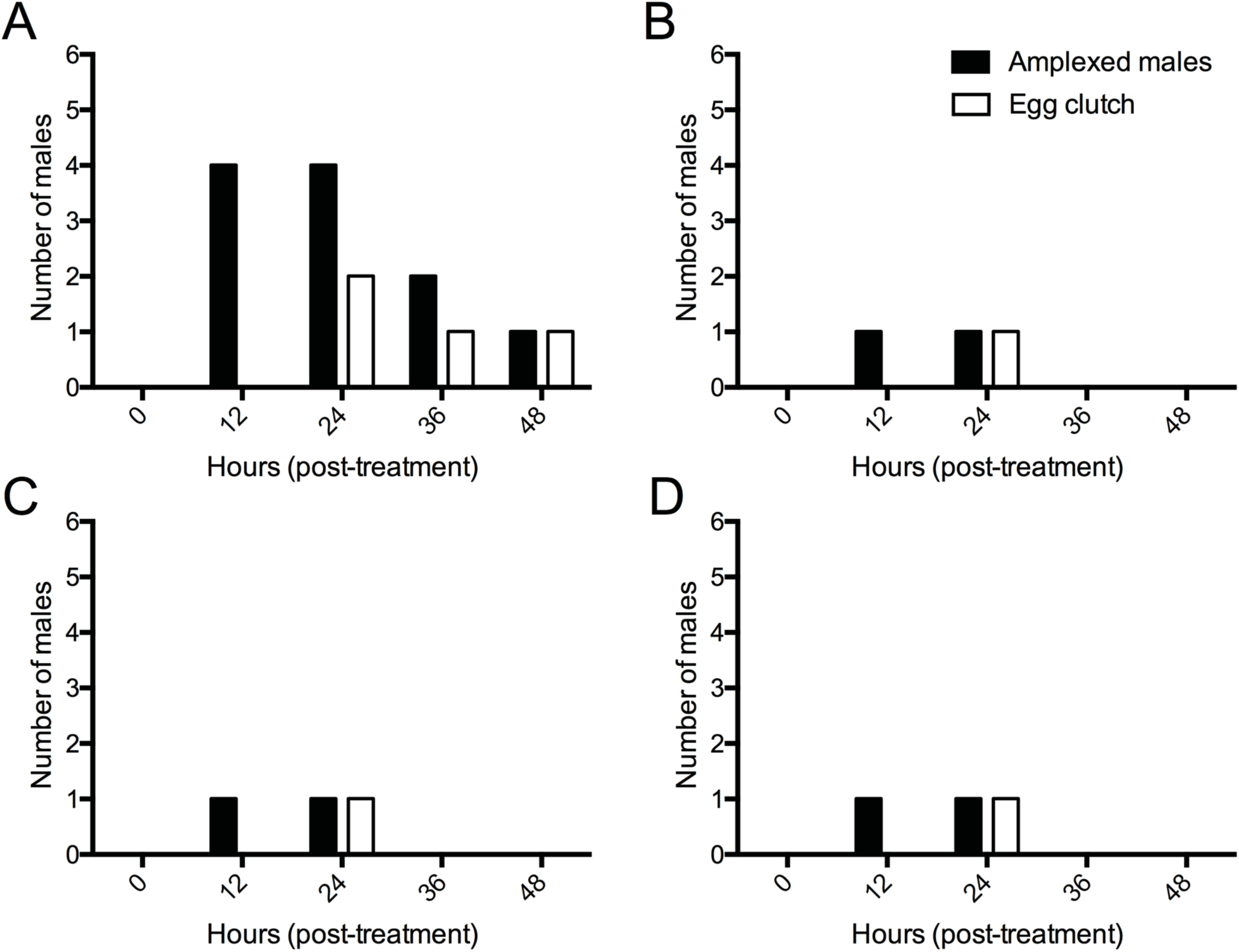
The total number of amplexed males in each experimental group (A–group A, B–group B, C–group C and D–controls) during 2013. The five time points shown (0, 12, 24, 36, 48 h) reflect the number of hours post–treatment that males were observed in amplexus and egg clutch represents the time post–male treatment at which egg masses were first detected. Only results for egg clutches deposited by amplexing pairs are represented.

## DISCUSSION

In captive amphibian breeding facilities, loss of reproductive capacity has often been associated with the absence of environmental cues found in the wild. In the absence of appropriate environmental conditions, previous studies of the boreal toad demonstrate that hormones can be administered to promote gamete release in males and females (Roth et al. 2010; Calatayud et al. 2015). In the current studies, we examined the effects of the previously reported hCG and GnRH protocol for females (Calatayud et al. 2015) and a single dose of hCG and GnRH for males (Roth et al. 2010) and their effects on the duration of amplexus, sperm and egg release.

Our results indicate that hormone administration significantly affected oviposition but did not significantly affect a treated male’s probability of amplexing. Since a greater number of treated females oviposited compared to controls, regardless of whether they had been amplexed, these results indicate hormones can induce oviposition independently of amplexus. Sexual receptivity in female anurans has been correlated with elevated gonadal steroids, readiness to oviposit and egg maturity (Wilczynski et al. 2005, review). Therefore, initial female resistance to a potential mate may decrease as circulating estrogen and progesterone reach peak levels (Lynch and Wilczynski 2006). Female receptivity would increase as egg maturation and ovulation was induced by hormone treatment and may provide a strong signal to nearby mates regardless of whether the males have received a hormone treatment.

As mentioned in our previous study, it is likely that priming doses of hCG caused a gradual rise in circulating LH levels in treated females, which culminated in an LH surge and increased progesterone production in developed oocytes, resulting in ovulation (Amsterdam et al. 1989; Fernandez and Ramos 2003; Calatayud et al. 2015). However, we suggest that amplexus with a male causes the female to oviposit faster.

Results from 2013 indicated that females that were amplexed by males that had been injected 12 h prior to the female receiving an OvD oviposited at significantly earlier times. Although amplexus did not significantly affect oviposition, the external stimulation provided by the male may provide a physical mechanism that increased the rate at which the eggs travelled through the oviduct, compared exclusively to the abdominal contractions observed in the females. Nevertheless, it is unclear whether the influence of amplexus on oviposition was overshadowed by the stronger influence of hormones, since we did observe oviposition by one control female after becoming amplexed.

In anurans such as *Bufo japonicus,* under normal conditions, amplexus causes an LH surge, which in turn stimulates spermiation (Ishii and Itoh 1992). GnRH stimulates the brain to produce LH and FSH in the pituitary and, as shown in *Bufo cognatus*, the testes produce sperm and the associated Leydig cells produce increased levels of testosterone (Propper and Dixon 1997). The induction of spermiation before amplexus in hormone treated toads may be the result of hCG acting directly on the testes before GnRH can act on the brain to elicit an LH surge. In this instance, amplexus may occur after spermiation because hCG stimulates protein secretion by Sertoli cells which, in turn stimulate Leydig cell steroid biosynthesis increasing testosterone and with it reproductive behaviors as reported in mice (Onoda et al. 1991; Langhorne 2016). While hCG may be directly affecting earlier spermiation in the testis, hCG may also enhance GnRH induced LH surge, which results in amplexus after spermiation. The detection of sperm at 24 and 48 h PT suggests that, once the behavior has been elicited, amplexus is sufficient to maintain the complete reproductive responses of the male. This theory supports our results, which show that males injected 12 h before females remained in amplexus for longer than males that were injected at the same time or 12 h after females. This is not surprising because males in group A would have to wait longer for the females they amplexed to be ready to oviposit, compared to males that were injected at the same time or 12 h after females.

In amphibians, the administration of hormones and their influence on egg maturation is not well understood, nor are the effects of repeated, long–term hormone treatment. Management should also consider the possibility that females may not breed every year and may skip two or more breeding seasons before ovipositing again (Carey et al. 2005; Muths et al. 2010). Additionally, we have not explored the effects of age, breeding history and frequency of hormone use on the reproductive health of these animals. Analyzing egg quality in females of different ages may be necessary to determine if breeding females over a certain age results in low fertilization rates or poorer quality offspring. Age assessments, with regards to male contribution to fertilization, should also be studied in the future.

This study demonstrated that, although exogenous hormones are not required to induce amplexus in the captive boreal toad, injecting males may; (1) be beneficial in increasing the length of time they amplex and spermiate and (2) increase the chances of synchronizing gamete deposition.

## CONCLUSIONS

It appears that it is not necessary to hormonally treat male boreal toads to induce amplexus or spermiation. However, males that are hormonally treated 12 h before females receive an OvD amplexed for longer, which implies that hormone treatment may be important for gamete synchronization in captive breeding efforts. It may also be favorable to expose males to females when they are receiving priming doses, to allow males’ time to respond to the female’s physiological response.

## Acknowledgements

This study was supported by an Institute for Museum and Library Services (IMLS) National Leadership Grant (LG–25–09–0064–09). The Native Aquatic Species Restoration Facility (NASRF) Alamosa, Colorado Animal management and research studies reported here were reviewed and approved by the Mississippi State University Institutional Animal Care and Use Committee (IACUC # 10–082). We thank our collaborators at the Colorado Division of Parks and Wildlife, H. Crockett and M. Nicholl as well as Mr. T. Smith Hatchery Manager and the staff at the Native Aquatic Species Restoration Facility in Alamosa, Colorado for allowing us to perform our studies. In particular, I thank Dr. S. Willard and Dr. A. Kouba for their suggestions and editing of the manuscript, Dr. C. Williams for statistical advice, Mr. T. Mix, Mr. J. Houghtaling, Mr. D. Westerman and Mr. N. Heredia for assisting me with my experiments during my 2012 and 2013 seasons at NASRF.

